# GeneDrive.jl: A decision tool to optimize vector-borne disease management planning under climate change

**DOI:** 10.1101/2024.08.23.609480

**Authors:** Váleri N. Vásquez, Erin A. Mordecai, David Anthoff

## Abstract

We introduce GeneDrive.jl, the first software package to optimize operational planning for the biological control of mosquito disease vectors. Mosquitoes are responsible for transmitting a significant percentage of the global infectious disease burden, a problem being exacerbated as climate change shifts the range and alters the abundance of these thermo-sensitive arthropods. But the efficacy and cost of vector control varies according to species, region, and intervention type. Meanwhile, existing computational tools lack the ability to explicitly tailor interventions for local health objectives and resource limitations. GeneDrive.jl addresses this equity and efficiency gap, which is of particular concern for the under-resourced nations that both bear the highest mosquito-borne disease burden and are subject to disproportionate climate impacts. The software customizes management strategies according to specific health goals and financial constraints, and can also be used to characterize risk by analyzing the temperature-responsive dynamics of wildtype vectors. GeneDrive.jl is designed to accommodate two important realities shaping the future of vector-borne disease: first, the genetic-based technologies that are defining a new era in control, and second, the uncertainty that increasingly variable and extreme temperatures bring for the climate-sensitive illnesses transmitted by mosquitoes. The software provides a ‘build once, solve twice’ feature wherein users may define a health management problem, optimize it, and subsequently subject outcomes to scenario-based testing within a single coherent platform. We demonstrate the policy relevance of this scalable open-source framework via case studies featuring *Aedes aegypti* in the dengue-endemic region of Nha Trang, Vietnam.

**Author Summary:** We present a software package designed to optimize and simulate genetic biocontrol, a broad suite of technologies that alter the genotype or phenotype of mosquito disease vectors by suppressing or wholly replacing vector populations. Our GeneDrive.jl library responds to a need for the fast, temperature-sensitive, low-cost exploration of public health management options, particularly in under-resourced global regions most at risk for current and future mosquito-borne illness under climate change. It is the first software to offer mathematical methods that optimally tune intervention strategies to local human health goals and resource limitations. Additional mathematical methods within GeneDrive.jl permit the simulation of optimized results given alternative parameterizations, furnishing a unique and scientifically important capacity to subject outcomes to scenario-based testing. The software is constructed to evolve along with the latest science, prioritizing composability^⊥^ and modularity^⊥^ to enable iterative updates without requiring a full rewrite. GeneDrive.jl addresses the confluence of two rapidly changing realities – existing and projected climate warming, together with advancements in biocontrol technology – when the state of the art, namely past field work and expert opinion, are no longer reliable guides for future planning. This paper is intended for an interdisciplinary audience and includes a Glossary to facilitate reading (see^⊥^).

## Introduction

Under-resourced nations and communities bear the brunt of mosquito-borne disease, a massive global burden that saw unprecedented surges in 2023 and 2024.^1^ The annual human health impact attributed to symptomatic dengue infection is 100 million people across approximately 125 countries, while treatment worldwide is estimated to cost $8.9 billion per year.^2–4^ Meanwhile, the year 2022 saw an estimated 249 million cases of malaria in 85 endemic countries; that disease carries a $12 billion annual pricetag in direct costs for Sub-Saharan Africa alone.^5,6^ Climate change now presents a compounding public health threat to this and other low-income regions of the world: thermal biology plays a central role in both vector population dynamics and pathogen infection and transmission from the vector.^7^ A warming world is expected to increase net and new exposures to the arboviruses transmitted by the *Aedes* mosquito, while some regions will see extensions of seasonal suitability for the *Anopheles* mosquitoes responsible for spreading malaria.^8–10^ By 2080, more than 60% of the global population is predicted to be at risk of dengue exposure.^11^ Such projections, together with the recent severe spike in disease incidence, illustrate the importance of proactive management.

Vector population control is an important means of mitigating mosquito-borne disease; however, costs vary widely according to technology and implementation, and evolved resistance is undermining the efficacy of conventional insecticide-based methods.^12^ Genetic-based innovations are recent alternatives to traditional prevention approaches, some have already been deployed in trials around the world.^13,14^ Genetic biocontrol alters the genotype or phenotype of a vector species through engineering, radiation, or deliberate bacterial infection.^15–19^ But the field work historically used to evaluate these methods is expensive, time consuming, and highly localized. Computational tools provide an affordable means to assess suitability for regions that may benefit from these and other technologies. Existing frameworks with varying mathematical approaches and degrees of specificity calculate the impact of environmental factors on disease vectors.^20^ Others additionally permit the simulation of genetic biocontrol.^21,22^ All currently available software stops short of including optimization methods to recommend mathematically feasible intervention^⊥^ strategies that satisfy user requirements.

GeneDrive.jl addresses this critical gap. It takes a significant step toward enhancing efficiency and equity in vector-borne disease management as the first domain-relevant library to include optimization capabilities, a feature that enables planners to predicate intervention decisions on regional resources and public health goals. GeneDrive.jl uses a temperature-responsive dynamic model^⊥^ that is comparable to previous efforts.^21^ The package’s primary innovation is its decision model^∞^, a constrained nonlinear program^∞^ (NLP) formulation of the temperature-responsive dynamic system (see Methods). Constraints^∞^ include biological, climatological, geographic, and budgetary factors that restrict the feasible solution space for a potential management plan. For example, thermal fluctuations that affect vector population size may dictate the ideal timing of a given intervention, the spatial layout of a neighborhood may influence the best location to conduct that intervention, or finances may limit available intervention resources.

Presently, software for vector-borne disease management allows the post-hoc evaluation of intervention plans, and can be used to conduct a parameter sweep approach to intervention design. This method involves running multiple consecutive iterations of a simulation to evaluate a range of potential outcomes and identify the best plan possible within the scope of that sensitivity analysis. GeneDrive.jl’s decision model inverts this process to output optimal policies given defined goals and conditions as inputs.

Important earlier work has used optimal control to quantify the intervention schedules needed to achieve specified reductions in mosquito vector populations.^23–25^ But these stand-alone academic analyses are not easily extended: a new problem must be formulated to accommodate each data update or alternative scenario. Obtaining such closed-form solutions also requires reducing or omitting modelled details of the ecological system,^26^ limiting the operational relevance of results.^27^ Beyond the innovation and benefit of providing a domain-specific software framework for optimization, GeneDrive.jl’s decision model employs numerical mathematical programming^∞^ rather than analytical optimal control ^∞^. This enables the inclusion of biologically detailed constraints and complex objective functions^∞^, allowing realistic and thus actionable results. It also permits scientists to rapidly, iteratively experiment with the space of possible outcomes because updates are straightforward and computationally inexpensive to re-run.

A second contribution of this package is that its decision model permits stochastic optimization^∞^ under environmental uncertainty: when employed in its stochastic formulation, GeneDrive.jl accounts for daily weather variability by accommodating multiple alternative temperature scenarios rather than just one average scenario. This is an important feature because characterizing climatic fluctuations in expectation (i.e., averages) omits the effect of extreme events. Such exclusion is a public health issue when climate science shows that environmental edge cases can be expected to increase in frequency and intensity, and may have disproportionate impacts on vectors and transmission. By explicitly considering a range of potential weather manifestations to obtain a single intervention schedule that collectively balances the potential population-level effects of various projected scenarios,^28^ stochastic optimization via GeneDrive.jl offers a novel and powerful tool for climate-aware intervention planning. Policies produced by the stochastic decision model are more robust and adaptive to temperature-driven uncertainty, a critical concern given the thermobiology of vector dynamics and the potential for intervention efficacy to be influenced by heat in a warming world.^29^

GeneDrive.jl’s dynamic model, which employs the same underlying population equations as its decision model, makes it feasible to explore prescribed intervention schedules under different scenarios (e.g., perturbed environmental regimes or alternative parameterizations of the biological system; see Fig. 1).^15,29,30^ This permits users to study downstream details of the planning problem. Moving beyond intervention design, the dynamic model also allows scientists to model baseline vector dynamics absent population control, or replicate empirical field deployments. This is useful for basic understanding of the study system. For example, it can help to characterize the behavior of interacting ecological effects. Like existing software options, the dynamic model may be used to conduct a parameter sweep^∞^ approach to management planning. Because the dynamic model is an ordinary differential equation (ODE) formulation of the decision model, and parameterized using the same structured data inputs (i.e., data model^∞^), it is straightforward to develop a GeneDrive.jl workflow wherein optimal intervention schedules are first defined using the decision model and then evaluated under alternative assumptions using the dynamic model (e.g., assumptions about environmental conditions, choice of intervention technology, the wildtype population’s dynamics, size, or spatial structure, etc. See Methods.).

**Figure 1:**
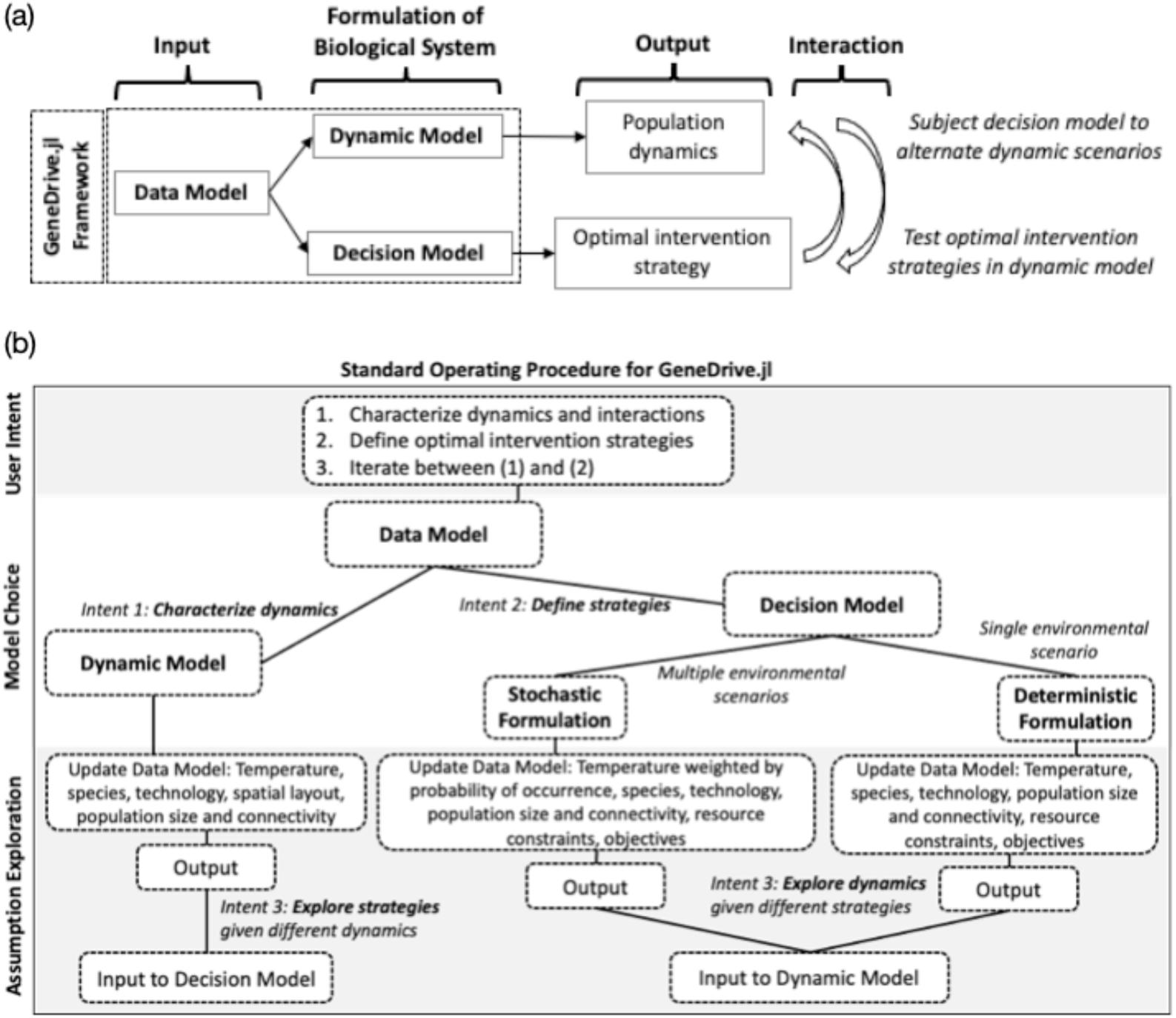
Graphical overview of and Standard Operating Procedure for the GeneDrive.jl package. Panel (a): GeneDrive.jl is a three-part framework (Data Model to standardize information inputs, Decision Model to optimize intervention strategy, and Dynamic Model to characterize ecological interactions and evaluate optimized management plans under alternative scenarios). It builds on the methodological innovations of Julia, the underlying programming language. Panel (b): Standard Operating Procedure (SOP) defined according to user intent, where the first step involves creation of the Data Model, the second step selects between the Decision and Dynamic Models, and in the third step users explore alternative parameterizations and iterate between the models as desired.

Researchers as well as public health decisionmakers seeking solutions require computational tools that are easily altered and scaled. This facility – a concept sometimes called composibility^∞^ in computer science – is essential for mathematical models to evolve with new technological discoveries and changing climatic realties. The methodological innovations offered by GeneDrive.jl build upon software design choices that prioritize composability and leverage inherent features of Julia, the software’s underlying programming language, to do so. For example, the GeneDrive.jl data model – a series of structures for standardizing information inputs – categorizes data and associates specific behavioral functions with those categories, also called Julia types.^31^ One application of this approach is that the species data already included in GeneDrive.jl reflect unique thermal responses when the user declares the appropriate category, requiring no further parameterization (presently *Aedes aegypti* and *Anopheles gambiae*; see Methods). Julia’s ecosystem of mature mathematical packages and large suite of free solvers gives GeneDrive.jl the ‘build once, solve twice’ workflow feature critical to policy design and evaluation: the dynamic model is constructed using the ODE methods in the DifferentialEquations.jl^32^ package while the decision model employs the optimization methods provided by JuMP.jl.^33^ An open source high-level language with C++ comparable speed, Julia also democratizes access to evidence-based planning by supplying up-to-date versions of all relevant dependencies^∞^ and recording compatibility constraints^∞^ via its package manager^∞^. This eases user experience of GeneDrive.jl by rendering a “ready-made” working environment^∞^.^34^ A visual summary of the software highlights key elements of its structure and operating procedure (Fig. 1).

To demonstrate GeneDrive.jl’s capabilities we present results that optimize risk reduction and cost savings goals using case studies of *Ae. aegypti*, a primary vector of dengue, in the region of Nha Trang City, Vietnam.^35^ Model outcomes accommodate historical observed daily temperature data for that location, as well as Coupled Intercomparison Model Project 5 (CMIP5) projections of future conditions under Representative Concentration Pathway (RCP) 8.5 scenarios (the most extreme greenhouse gas concentration trajectories adopted by the Intergovernmental Panel on Climate Change (IPCC)). Each example employs a genetic biocontrol method called Release of Insects carrying a Dominant Lethal gene (RIDL), an engineered suppression technology that eliminates competent vector populations and has been the subject of multiple field trials. We compare solutions derived using the deterministic and stochastic iterations of the decision model, and evaluate the human health and economic gains made possible by accounting for locally-specific details including daily weather variability. We also demonstrate the iterative workflow possible between the dynamic and decision models, which is conducive to adaptive policy development. Each example advances transparency by illustrating how to build a GeneDrive.jl model, input user-specific datasets, and take advantage of the sample parameters stored in pre-defined data model files.

## Results

### Model overview

Optimized intervention policies are produced by a nonlinear program that schedules the location and frequency of biocontrol deployments, along with the number of transgenic organisms that comprise each release. This program is formulated first as a deterministic and then as a stochastic model, where the former takes daily weather inputs as a single time series of temperature and the latter takes a matrix of multiple time series, each of which is weighted by the probability of its occurance. Initially all temperature time series in the case study are assigned equal weight to illustrate the outcome of stochastic optimization. We then demonstrate how to explore specific temperature scenarios happening with higher likelihood, illustrating the notable policy benefits of this feature. A diagram of the decision model information flow illustrates the difference between stochastic and deterministic inputs (Figure 2).

**Figure 2:**
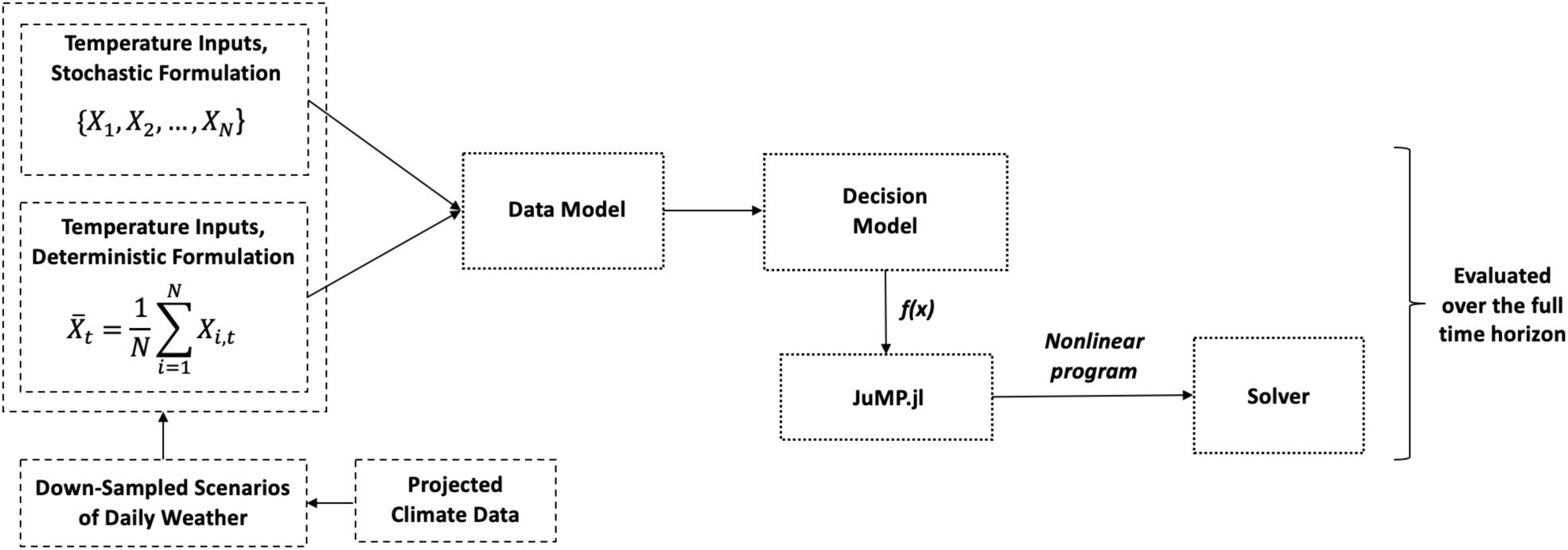
Decision Model information flow, where the set {X_1_, X_2_, …, X_N_} contains N timeseries of daily temperature and \bar{X}_t_ reflects a single time series of daily temperature t, which may be an average over N time series of daily temperature X_t_. An example source of temperature information is daily weather time series down-sampled from scenarios of climate projections. Once parameterized via inputs to the Data Model, the Decision Model function f(x) is passed to JuMP.jl solvers.

Additional case study parameterizations input via the data model include details of vector species, biocontrol technology, and the geographic topology across which the intervention is being optimized (see Methods). Optimization is performed by maximizing or minimizing a customizable objective function (see Methods). All examples here employ the same goal, also called an objective function (Equation 1). Eqution 1 jointly minimizes the vector competent (wild adult female) population *F_g_* and the number of modifed adult male mosquitoes *c_ĝ_* released to mitigate them. Variables *ĝ* and *g* are indices of the set of genes *G*, respectively indicating modified and wildtype genotypes. The minimization is sought in each daily time step *t* by a factor of 20% (*ψ* = 0.20) with respect to the size of the population on the first day of the simulation (*t* = 1), from a designated date (*t* = 200) through the problem horizon *T* (*t* = 365). While this work showcases a year-long period, the model may be applied to shorter or longer time frames of interest.

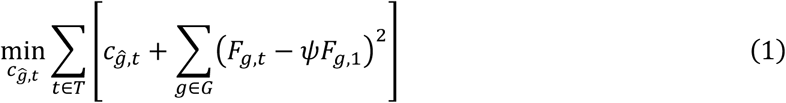

### Decision model: Deterministic optimization

From a planning perspective, deterministic constrained optimization facilitates the design of locally tailored intervention strategies. It explores the solution space comprehensively, unlike the more widely used process of manually evaluating individually crafted policies. Optimization reduces computational effort compared to the current state of the art and assists in discovering potentially unintuitive solutions. Here, we show its application as a tool for achieving public health goals in a maximially resource- efficient manner given limitations imposed on the frequency of field deployments. These limitations represent constraints on site accessibility, budget, or available labor. We then demonstrate the model’s utility as climate change alters the conditions under which interventions are deployed, undermining historical precedents of what works. The flexibility to quickly and cheaply optimize for shifting scenarios improves policymakers’ ability to adapt decisions even as new economic, ecological, or technological realities emerge.

A series of deterministic decision model runs show the impact that the hotter and more variable environmental conditions may have on optimal intervention schedules (Fig. 3). Optimizations are conducted using a time series of average observed temperatures from the Nha Trang City region of Vietnam during the 2000s as well as an average of time series projected for the 2030s. The resulting policies produced by the model differ both in the total number of modified organisms used and prescribed trips to the field. We compare outputs when the solution space is constrained to weekly versus biweekly release frequencies under historical and future temperature regimes, respectively.

**Figure 3:**
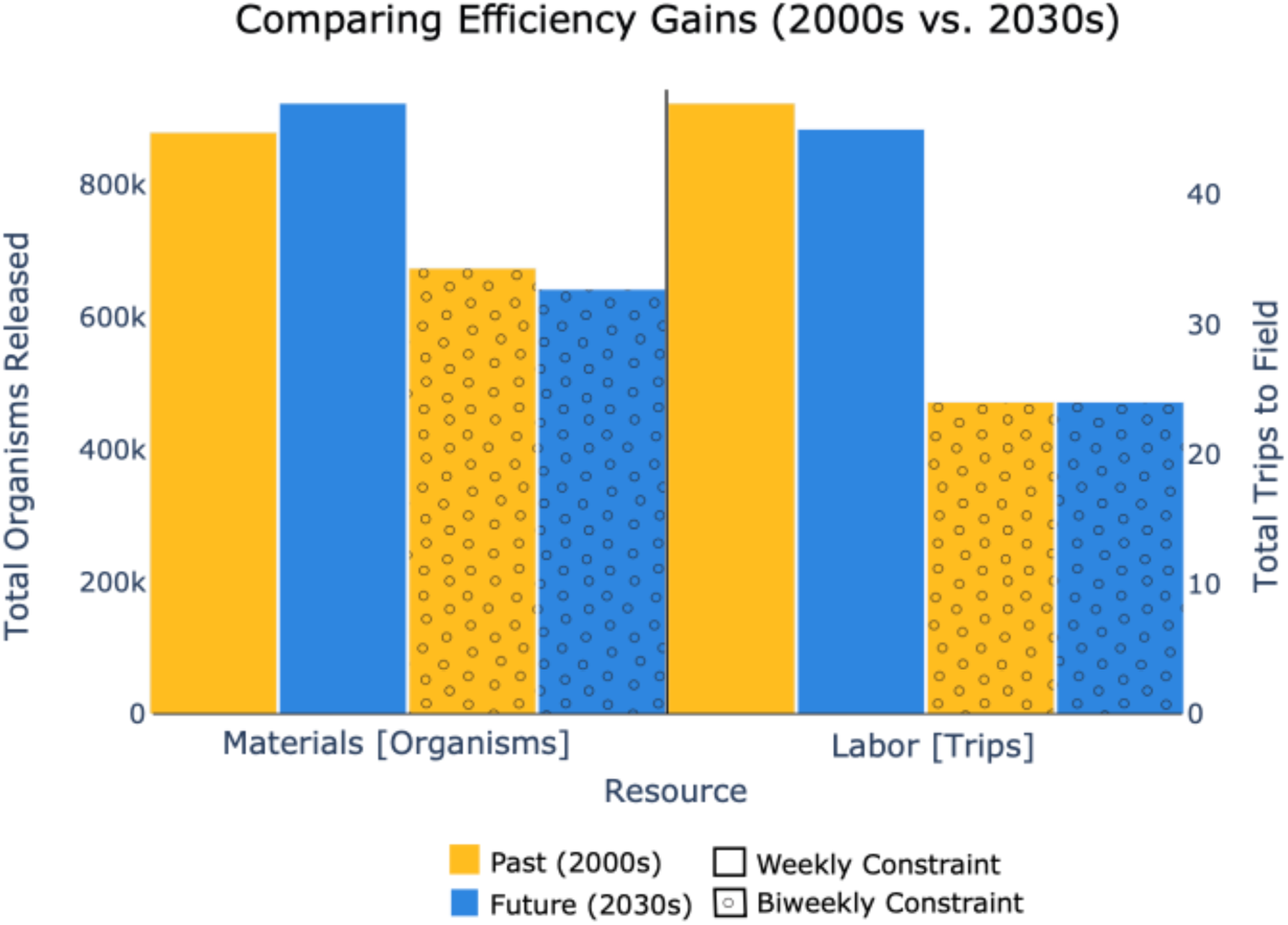
Optimized weekly and bi-weekly deployment policies under historic and future temperature regimes show that bi-weekly strategies allow for resource conservation^∞^ in both cases while still achieving the stated public health goal. Results show that optimization is useful for exploring solution space: it can accommodate necessary constraints (e.g., deployment frequency) without compromising risk reduction. Historic scenarios using bi-weekly strategies save comparably more on trips to the field, while future scenarios using bi-weekly strategies save on material costs (total transgenic organisms released). The difference in resource expenditure (materials and labor) for prescribed weekly (solid colors) and biweekly (dots) schedules under both historic (yellow) versus future (blue) regimes is shown.

Across both historic and future schedules, we observe efficiency gains when imposing biweekly operational limitations. The decrease in resource expenditure seen when applying biweekly constraints to the historic scenario (a decadal average of annual observed temperature timeseries in the 2000s) amounts to a difference of 205,121 total organisms and 23 individual deployments. In the policies optimized for the future scenario (a decadal average of annual projected temperature timeseries in the 2030s), 281,097 fewer transgenic mosquitoes and 21 fewer release instances are employed for the biweekly strategies as compared to the weekly. Under the weekly constraints, projected climate change will require more transgenic mosquitoes but fewer field site visits compared to historic temperatures. The discrepancy amounts to an additional 44,171 organisms and a reduction of two discrete deployments – from 47 to 45 trips – compared to the past. Given the biweekly constraint, 31,805 fewer mosquitoes would need to be released under future projected temperatures, but the same number of trips to the field is called for in both historic and future scenarios.

### Decision model: Stochastic optimization

From the perspective of resource savings for real-world practicioners, the stochastic formulation of the constrained optimization problem applied to the Vietnam example demonstrates significant advantages over the deterministic approach. We show that those material gains stand to increase with the hotter and more variable temperatures brought by climate change. The reason for this is both biological and methodological. First, as environmental fluctuations grow more extreme and those extremes become more frequent there will be greater variability in the vector competent population; this demands management that is sensitive to such dynamics. Deterministic optimization is capable of accommodating variability for one projected scenario of annual temperature at a time. However, because weather predictions carry uncertainty, it is more realistic to recognize a range of potential futures. Doing so with the deterministic model implies averaging the various potential scenarios, a characterization which would smooth the appearance of extremes. But the stochastic formulation permits consideration of multiple temperature time series in tandem, accounting for the potential outliers of each uncertain projection and returning an optimized policy that balances the possibility of their occurance.

As shown here, the stochastic decision model yields biocontrol strategies that employ hundreds of thousands fewer modified organisms across both historic and future scenarios (Fig. 4). Policies produced by the stochastic program reduce trips to the field over the course of both historic and future modelled years by approximately 30% compared to deterministic optimization. The value of accounting for the daily weather uncertainty of six observed and projected time series of annual temperature in the Nha Trang City region is illustrated by comparing stochastic decision model outputs (darker colors) to deterministic decision model results (lighter colors) for each regime (Fig. 4). Schedules are optimized for the historic (2000s) and future (2030s) period (Figs. 4a and 4b, respectively). When assessing the prescribed change in total numbers of released organisms, savings enabled by the stochastic compared to the deterministic model are 284% in the future case study, compared to 45% in the historical example.

**Figure 4:**
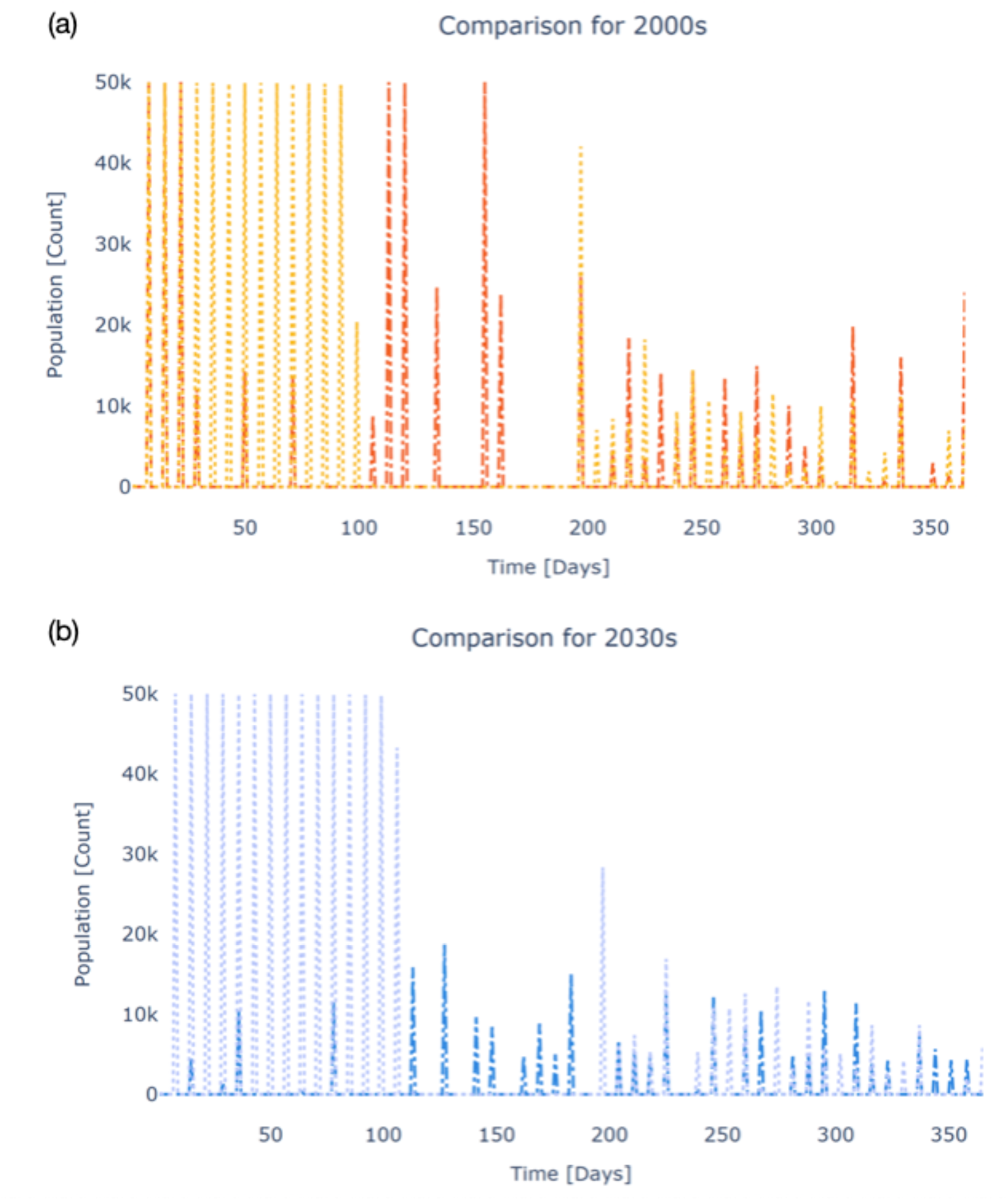
Comparing deterministic and stochastic optimized policies under historic (Panel (a), 2000s) and future (Panel (b), 2030s) temperature regimes shows that cost savings are significantly larger when applying the stochastic decision model to define future strategies. These gains are demonstrated both in amount of material used and total trips to the field. Panel (a) shows deterministic (yellow) and stochastic (orange) schedules given historic temperatures. Panel (b) shows deterministic (light blue) and stochastic (dark blue) schedules given future temperatures.

Thirty-two deployments are prescribed by the stochastic model under both historic and projected temperatures, compared to the deterministic model policy of forty-seven (historic) and forty-five (future).

### Dynamic model: Characterizing outcomes

After defining optimal intervention strategies using the decision model, we use the dynamic model of GeneDrive.jl to simulate those prescribed strategies and examine their effect on mosquito populations over time under different climate scenarios. This illustrates the tradeoffs of alternative policy options. We iteratively apply deployment schedules that have been deterministically or stochastically optimized to wild *Ae. aegypti* populations whose dynamics are responsive to four different temperature scenarios: two with high daily thermal variability and two with median daily thermal variability (see Methods). Results highlight that as temperatures become hotter and more variable, it will be increasingly important to analyze potential compromises between the economic cost and feasible health outcomes of vector-borne disease interventions, because these tradeoffs may become more stark.

Under historical temperature regimes, optimized deployment schedules – whether produced via stochastic or deterministic optimization – reduce the *Ae. aegypti* population to comparable levels (Fig. 5a). High versus median temperature variability scenarios do not produce significantly different results in this historical case study. When employing the stochastically-generated intervention policy, high variability scenarios briefly lead to elevated risk levels (a larger vector competent population). More heavily weighting the probability of the temperature scenario featuring the highest daily variability (see Methods) does not generate dynamics that are distinct from when all scenarios are weighted equally. From a health policy perspective these results suggest that the deterministic strategies may be a pragmatic choice assuming historical temperature scenarios. Stated resource constraints (material quantity and frequency of trips to the field) are met while achieving greater immediate health outcomes in the short term and approximately equivalent outcomes in the longer term compared to stochastic strategies.

**Figure 5:**
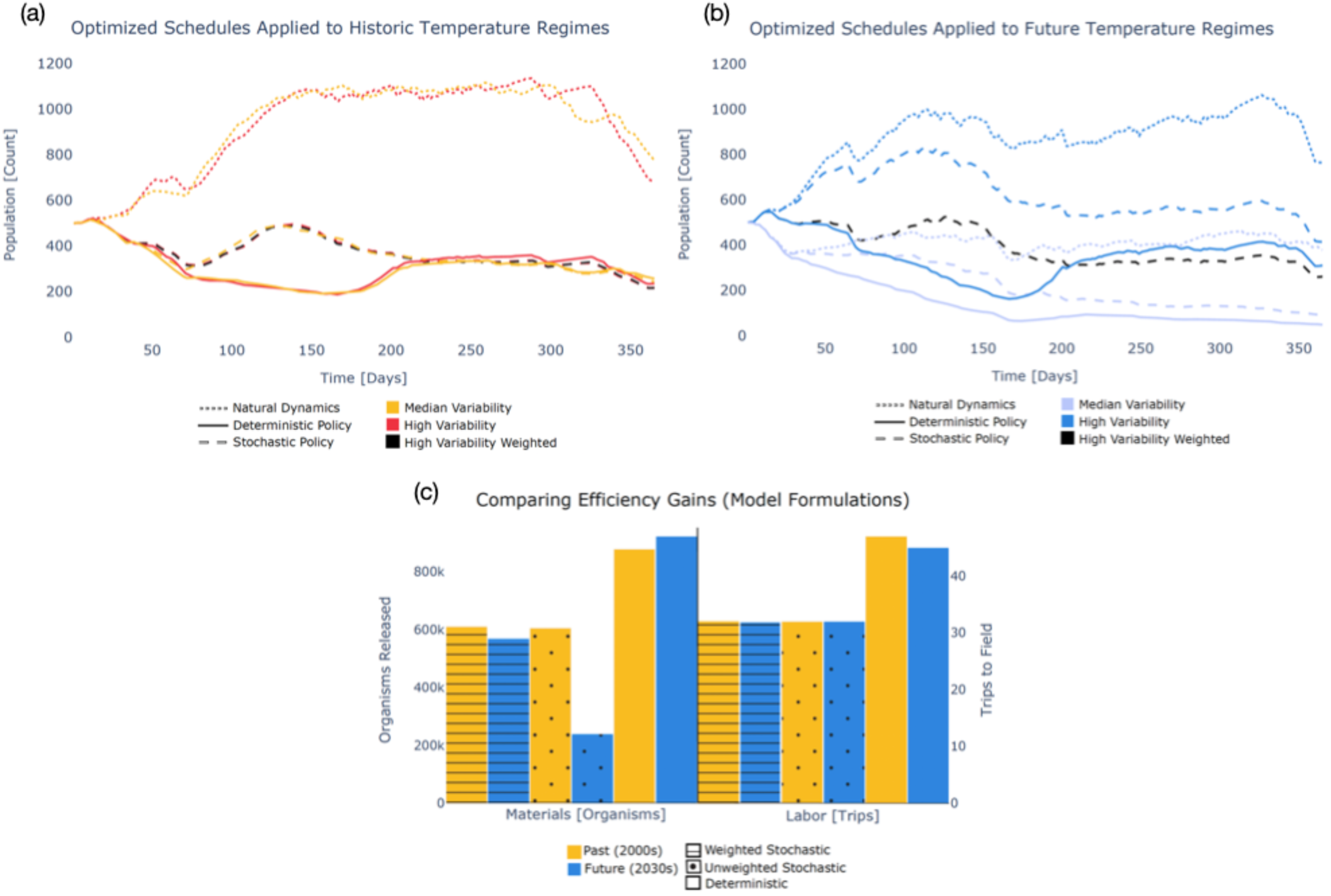
Population dynamics subjected to alternative intervention policies demonstrate the public health benefits of deterministic strategies under historic temperature scenarios as well as under future scenarios with median temperature variability. The comparable health benefits and cost savings of probabilistically weighted stochastic strategies is evident under scenarios of high future temperature variability. Panel (a) shows the suppression in natural vector population dynamics (dotted lines) resulting from policies determined by stochastic (dashed lines) and deterministic (solid lines) optimization, where yellow indicates median temperature variability, orange indicates high temperature variability, and black highlights the outcome of the weighted stochastic policy under the high variability scenario. Panel (b) again shows the suppression in natural vector population dynamics (dotted lines) resulting from stochastic (dashed lines) and deterministic (solid lines) policies, where light blue indicates median temperature variability, dark blue indicates high temperature variability, and black highlights the outcome of the weighted stochastic policy under the high variability scenario. Panel (c) shows deterministic (solid colors), probabilistically weighted stochastic (lines), and unweighted stochastic (dots) resource expenditure given past (yellow) and future (blue) temperature scenarios.

In contrast to the historical temperature regime, under future temperature projections with median daily variability, deterministically-defined interventions are 46.9% more effective than unweighted stochastic strategies in suppressing the vector population by the end of the observed period, but at a much higher resource cost (Fig. 5b). These results exemplify how the dynamic model furnishes policymakers with additional decisionmaking context: there is a bigger price tag incurred by the more resource-demanding deterministic release schedule; however, as evidenced by the dynamic model results, there are greater potential human health and consequent downstream economic costs stemming from the unweighted stochastic release schedule because a larger disease-competent mosquito population remains. When the future scenario with the highest variability is probabilistically weighted (black dashed line in Fig. 5b) to reflect the assumption – supported by climate science – that higher variation in daily temperatures will become more likely in the future (see Methods), the human health risk at the end of the simulated year is 18.9% lower than under the deterministic model strategy (dark blue solid line in Fig. 5b). Decisionmakers must consider such gains while taking into account the resource expenditures prescribed by the range of model formulations (Fig. 5c).

**Table 1:** The iterative policy relevance of each software feature in GeneDrive.jl. Columns highlight select package capabilities, explaining the application of each facet to policy practitioners. Rows denote whether the feature is found in the Data, Dynamic, or Decision models.

| Model | Extensible and scalable | Simulates time-varying behavior | Optimizes for constraints and objectives | Optimizes accounting for daily weather uncertainty | Policy Relevance |
| --- | --- | --- | --- | --- | --- |
| Data | **X** |  |  |  | Enhances transparency: model once, solve twice. Broadens applicability: modular updates as needed. Enables replication: stores inputs and relationships. |
| Dynamic | **X** | **X** |  |  | Facilitates sensitivity analyses: examine the effect of altered parameters. Allows scenario analysis: evaluate intervention impacts and tradeoffs. |
| Decision (Deterministic) | **X** | **X\*** | **X** |  | Permits context-relevant decisionmaking: define custom objective function as well as biological, budgetary, geographic, and climatological constraints. |
| Decision (Stochastic) | **X** | **X\*** | **X** | **X** | Accommodates uncertainty: optimize over a range of possible climates, probabilistically weighing the likelihood of each scenario. |

*\*When no objective is defined, model dynamics are not optimized. Rather, behavior is equivalent to an ODE simulation.*

## Discussion

Uncertainty in daily weather already presents a challenge for the design and implementation of biocontrol for public health. As temperatures warm and become more variable, the complexities inherent to mitigating mosquito-borne diseases – including the dengue virus explored in our case study – will continue to grow. For example, the field trials upon which current intervention planning are largely based will provide progressively less reliable guidance for tuning future deployment schedules. Vector-borne disease management thus stands to gain from computational methods, including the simulation approaches already in use, which incorporate mechanistic knowledge based on empirical experiments.

Here, we demonstrate the additional value of constrained optimization for public health purposes, and characterize the relative benefits of deterministic and stochastic decision models for policy applications. This work is directly responsive to the expert-expressed need for software decision tools that facilitate operational planning for climate-sensitive diseases.^20^

The hyper-local nature of mosquito-borne disease is driven by socioeconomic, environmental, and biological factors. To develop interventions that are regionally appropriate, these constraints must be taken into account explicitly. As demonstrated by multiple industries, including within the broader health domain,^36–38^ optimization via mathematical programming is a means of doing this while enabling the straightforward, iterative examination of alternative policy goals. We show that a deterministic nonlinear program produces health-effective and cost-efficient risk reduction assuming historical scenarios of daily temperature in the region of Nha Trang City, Vietnam, and that under the same historical climatic conditions a stochastic nonlinear program achieves nearly equivalent results while expending fewer resources in both labor and raw materials.

The intensity and frequency of heatwaves have been found to negatively impact the performance of at least one genetic-based intervention method,^29^ further highlighting the need for planning models that account for temperature-dependent dimensions of uncertainty. The results of our stochastic decision model illustrate that assigning greater likelihood to thermal scenarios that exhibit high daily variability produce intervention policies that reduce public health risks more effectively than those that treat all projected scenarios equally. While the examples presented here incorporate a critical exogenous stochasticity (i.e., temperature variability), it is also important to consider uncertainties additional to temperature, such as precipitation.^39^ Future work will expand the GeneDrive.jl model to accommodate more extensive uncertainty quantification, including with respect to the mechanistic modeling of vector species and genetic-based intervention technologies. While some of this work is underway, advanced users and those who seek to contribute to the development of GeneDrive.jl may leverage its modular design to extend or add to existing capabilities (see Methods).

Our dynamic model results showcase how daily temperature, including characteristics such as median versus high thermal variability, affects vector population dynamics. This exemplifies the utility, unique to GeneDrive.jl, of including a dynamic and decision model in a single coherent package: together, they facilitate a more comprehensive understanding of the conditions that contribute to differences in prescribed interventions as well as the potential range of outcomes from those policies. This can inform a richer evidence base for the discussion of optimized biocontrol. However, bridging the divides that presently exist between model outcomes and practitioner discussion requires developing more robust pipelines for collaboration.^40^ The roadmap that enables simulation results to guide policy implementation may differ regionally, and while recent efforts have begun establishing communities of practice to address this challenge,^20^ much work remains.

One step toward enhancing the policy relevance of software such as GeneDrive.jl is using it to design and evaluate a future field trial, such that results can be assessed using real-time empirical data to evaluate the potential effect of optimized schedules informing adaptive deployments. Another avenue for increasing the accessibility of software tools and allowing form to suit necessary function is to develop useability studies that include bench and field scientists as well as policymakers. This would challenge the current paradigm of models being largely academic exercises. The need for improved participatory approaches is not, however, limited to collaborations between modelers, scientists, and decisionmakers: community consultation is also a key component of ensuring that software design answers non-expert concerns, examples of which might include producing estimates of intervention technology risk or options to simulate and optimize remediation^⊥^ of genetic biocontrol.

## Box 1

**Key questions to which GeneDrive.jl might be applied.** Select examples of research or practice-oriented questions that could be addressed using the current iteration of the software include:

- Dynamic Model: Characterize the seasonal population fluctuations of mosquito disease vectors (e.g., *Aedes aegypti, Anopheles gambiae*) absent intervention, given alternative thermal scenarios.

- For example, how might specific species dynamics change under future temperature projections for Peru? For different regions of Brazil?
- Decision Model: Design locally targeted vector population management plans to address ongoing outbreaks or prevent minimize future disease risk while remaining within a defined budget (e.g., materials, labor, schedule).

- For example, what control strategy is optimal for the city of Singapore, given its unique goals and resources? How does this strategy compare to that recommended for the city of Dhaka, with its distinct geographic features, goals, and resource availability?
- Iterating between the Dynamic and Decision models: Compare the efficacy of technological interventions (e.g., suppression approaches using Release of Insects carrying a Dominant Lethal gene (RIDL) or *Wolbachia* infection, replacement approaches using Homing Gene Drive (HGD) or *Wolbachia* infection) under different temperatures, spatial topographies, and vector population size.

- For example, does network connectivity inform whether RIDL or Wolbachia succeeds sooner in reducing vector populations? Is effectiveness impacted by a tropical versus a temperate climate?

## ^⊥^Glossary

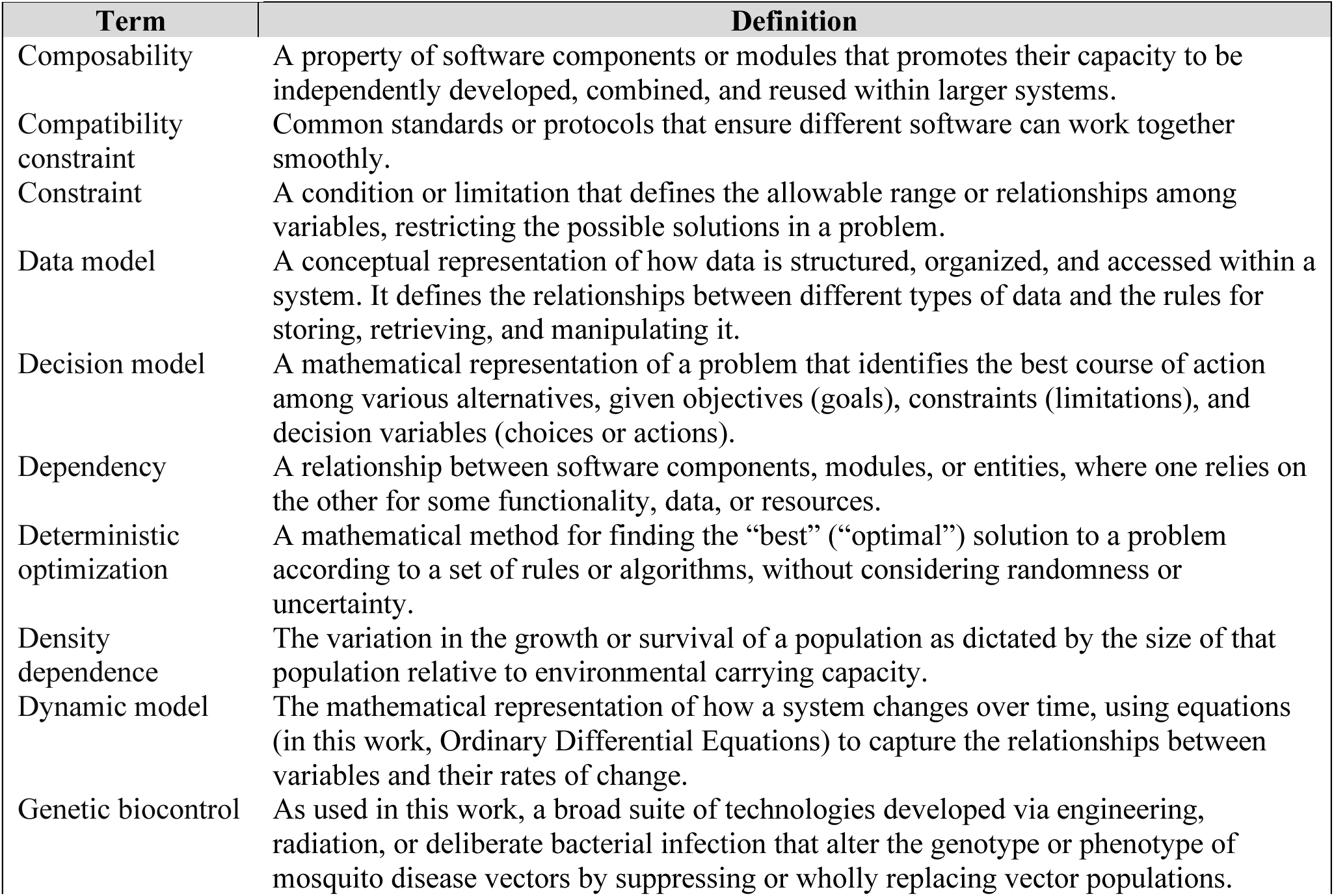
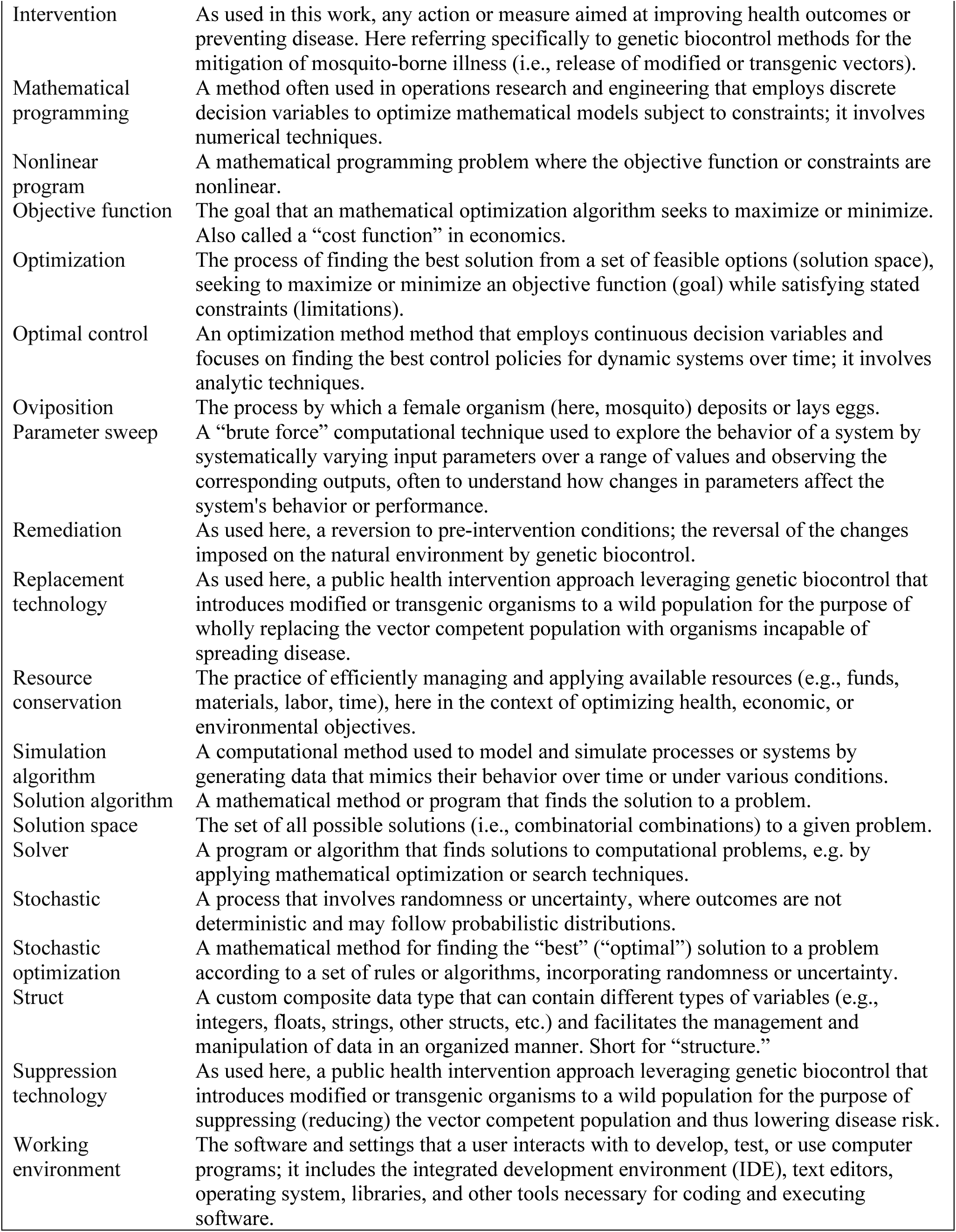

## Acknowledgments

The authors thank L. Wang for the data characterizing future heatwaves in Vietnam and guidance concerning these projections. The funders had no role in the study design, data collection and analysis, decision to publish, or preparation of the manuscript.

## Author contributions

Conceived and designed the paper: VNV. Collected the data, performed the analysis, wrote the first draft: VNV. Verified the data and analysis: VNV, DA. Revised the manuscript and approved the final draft: VNV, EAM, DA.

## Methods

### Section 1: Climate data

Historical daily temperature records for the region of Nha Trang City (Nha Trang), Vietnam (latitude 12.2534, longitude 109.1871) are from the Global Historical Climatology Network (GHCN) database maintained by the National Centers for Environmental Information of the United States National Oceanic and Atmospheric Administration (NCEI-NOAA). Two villages in the Nha Trang region, Vinh Luong Ward (Vinh Luong) and Tri Nguyen Village (Tri Nguyen), are the sites of recent genetic-based interventions for *Ae. aegypti*. Lacking weather data from the villages themselves, subsequent analysis of deployments used daily records from Nha Trang.^35^ Future heatwave data under Representative Concentration Pathway (RCP) scenario 8.5 was provided by Dong et. al. (2021).^41^ These data were used to develop projected heatwave scenarios for Vásquez et. al. (2023) using historical temperature data for the baseline period (1990-2005) as well as Coupled Model Intercomparison Project Phase 5 (CMIP5) projections of 2030 and 2050 temperature deltas for Vietnam from the World Bank Climate Change Knowledge Portal.^42^ See Vásquez et. al. (2023) for a thorough description of the anomaly method used to create the heatwave scenarios.

### Section 2: Data model

The data model is designed to compose each GeneDrive.jl problem using broad thematic components (e.g., temperature data, organism data). It stores the details of interest, enforcing consistency in the specification of data across computational experiments to facilitate both reproducibility and data sharing. ^43,44^ Each thematic component includes details that are themselves encapsulated in subcomponents (Methods Figure 1). This modularity promotes code re-use as different research questions or focal areas are studied; the functional form or parameterization of each component may be independently altered per problem specification without changing the larger code base.^45^ These structures grouping related variables, which can include different data types such as integers, floats, and strings, are also called “structs.”

Experimentation with different environmental assumptions is conducted by parameterizing one of three ‘Temperaturè structures. These include ‘ConstantTemperaturè to specify a static thermal environment, ‘SinusoidalTemperaturè for an idealized, seasonally fluctuating regime, and ‘TimeSeriesTemperaturè in which vectors of daily values can be stored. Each of these descriptions of thermal trend can be accentuated with heatwaves and cold snaps using the ‘TemperatureShockDatà structure to impose time-bound increases or decreases in temperature.

The ‘Organism’ type in GeneDrive.jl includes a ‘LifeStages’ data container for details of species-specific development time and mortality rate for juvenile and adult mosquito life stages. For select species (presently the primary vectors of dengue and malaria, respectively: *Ae. aegypti* and *An. gambiae*), empirically derived functions are included to characterize vital rate responses to environmental perturbations such as temperature. Static parameter values may also be used. The ‘Genetics’ data type within the ‘Organism’ component defines the likelihood with which offspring will be produced, as well as fertility rates, sex ratios, and varied fitness costs (e.g., the degree to which fecundity for modified organisms is biased with respect to their wild counterparts). It denotes the properties of the biocontrol technology being deployed (presently, options include Release of Insects carrying a Dominant Lethal (RIDL), *Wolbachia*, and Homing Gene Drive (HGD)).

For geographic information, the GeneDrive.jl data model provides a ‘Nodè and a ‘Network’ structure. The ‘Nodè datatype includes fields to store the ‘Organism’ and ‘Temperaturè components as well as geographic coordinates and location name. A ‘Network’ structure is comprised of a collection of ‘Nodès. Each species, life stage, genotype in a ‘Nodè is assigned a rate of dispersal using a transition matrix. Nonzero rate values instantiate bidirectional ‘Nodè connections. This feature enables users to account for a diversity of demographic and genetic migration tendencies as well as exogenous factors that may transport organisms from location to location.^46–48^

Anthropogenic actions such as biological control may be studied in the GeneDrive.jl dynamic model using the ‘Releasè object. This data type defines the time at which a certain number of organisms are added to or removed from a population during dynamic simulation. When specifying an intervention schedule, fixed or variably sized releases can occur at flexible intervals. Alternatively, interventions may be conducted in an adaptive manner that directly accounts for the size of the standing wildtype population using ‘ProportionalReleasè, wherein the release size is defined according to its relative magnitude with respect to the population of interest at a given timestep (e.g., 15% of wild females present on day 100 of the simulation).

For computational experiments that optimize anthropogenic interventions, the ‘ReleaseStrategy’ structure stores the information about operational requirements and limitations that is used in the decision model. The structure name is a nod to the biocontrol intervention strategies commonly used in the public health, agriculture, and invasive species arenas wherein organisms modified for, for example, sterility are released into the environment to reduce the standing wildtype population of a given species.^49^ The values of this structure are assigned on a per-node basis in the case of a network implementation, enabling spatially explicit constraints and the exploration of myriad policies. Default values are supplied where no information is specified; this information is accessible by viewing the fields of the data model.

**Methods Figure 1:**
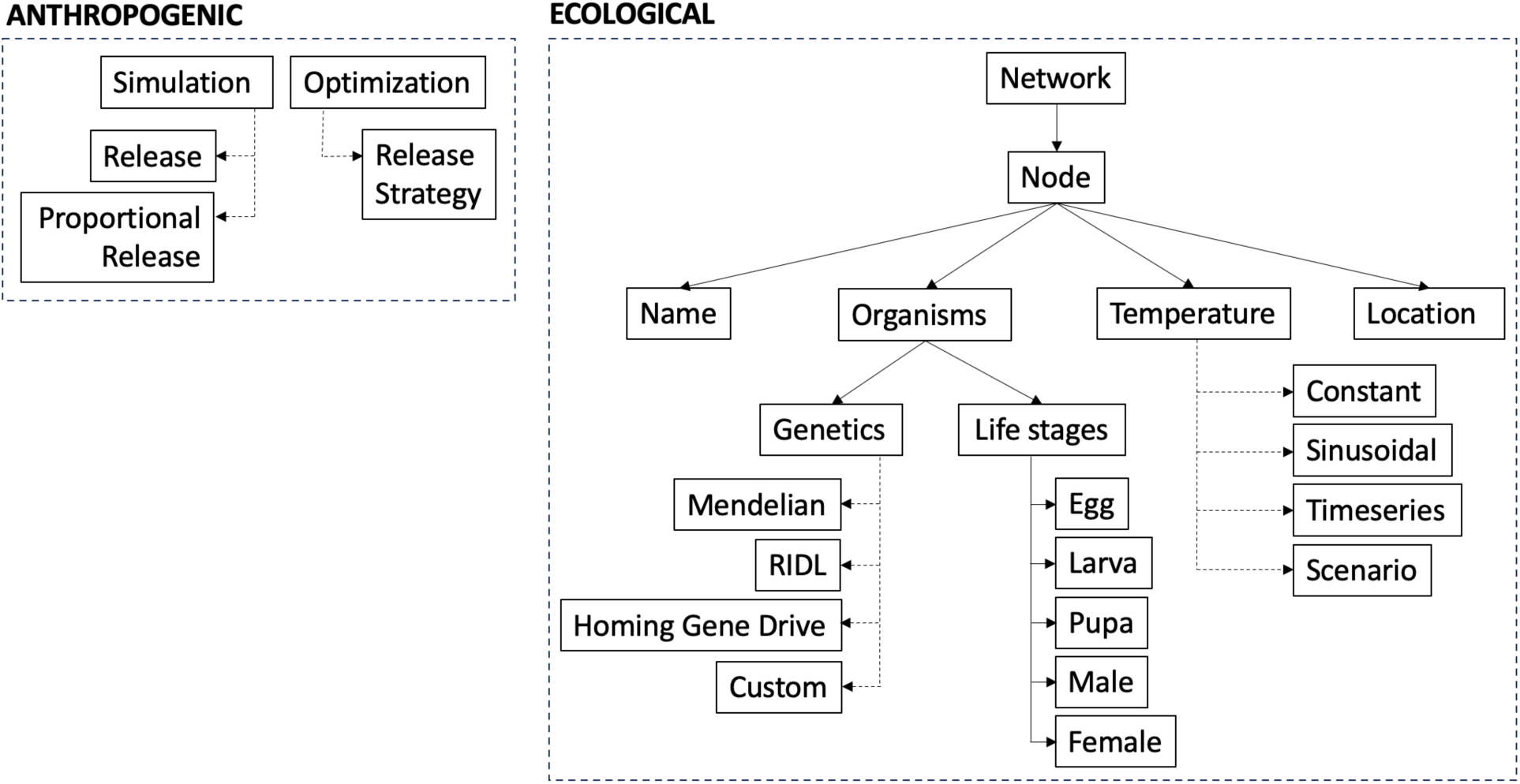
Abstract tree hierarchy illustrating the structure of data model components. Arrows with dotted lines indicate optional selections, and those with solid lines indicate components that are required to fully specify and store or run a GeneDrive.jl problem.

The case study presented in this work illustrates construction of the GeneDrive.jl data model (Code Block 1). It defines the vital rates, thermal biology, genetic characteristics, and habitat of an *Ae. aegypti* mosquito. The data model in this example is largely built by drawing from pre-constructed information stored in the GeneDrive.jl library. This highlights the code brevity and simplicity made possible by the software’s design, particularly once problem information has been assembled and saved. All data used in this case study are available in the package for replication and extension.

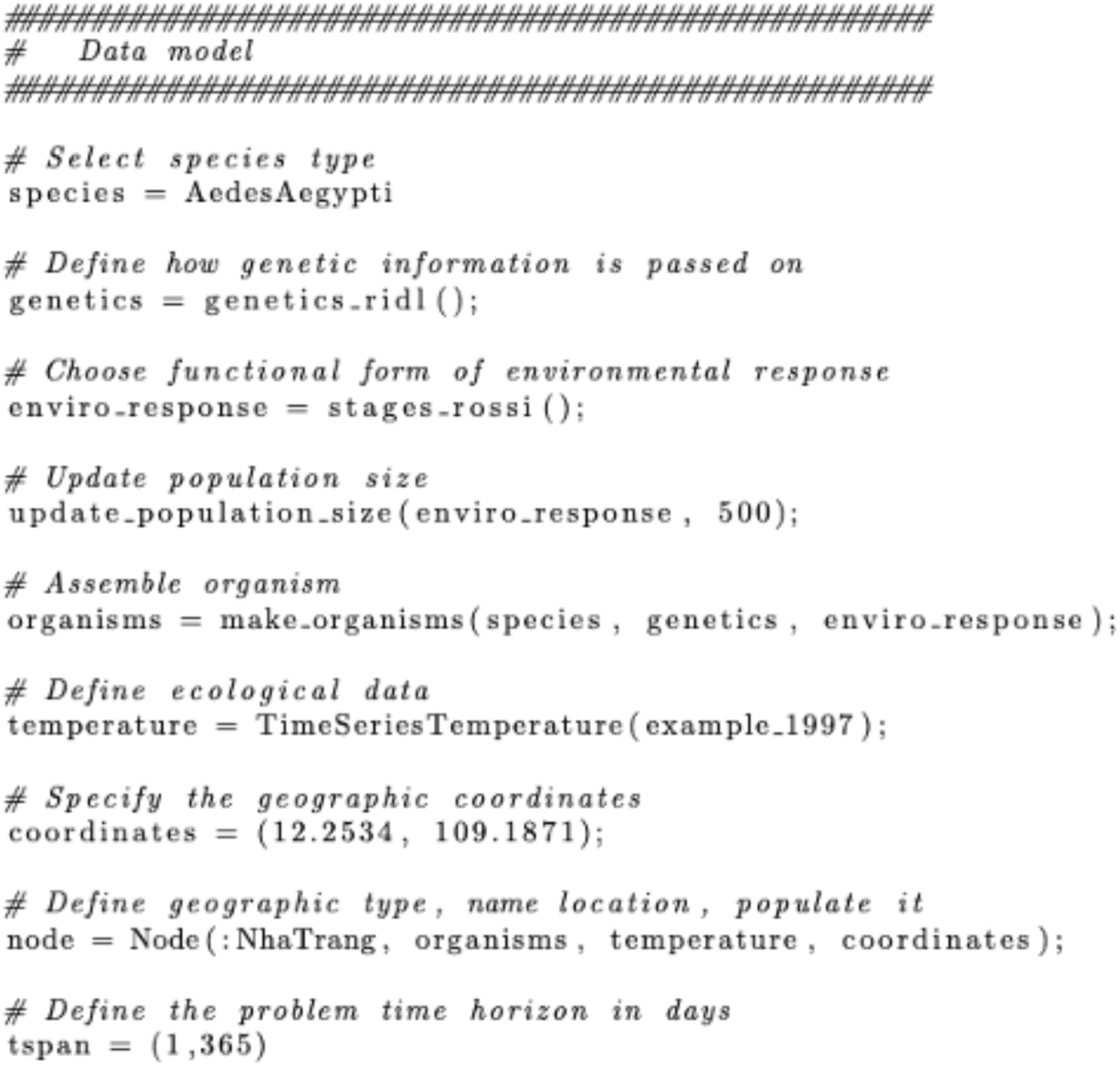
***Code Block 1:*** *Creating the data model.*

### Section 3: Dynamic model

The dynamic model in the GeneDrive.jl framework is a system of ordinary differential equations (ODEs) that simulates mosquito population dynamics (Eq. 1).

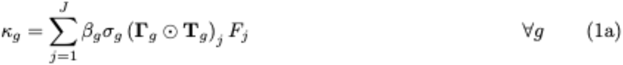

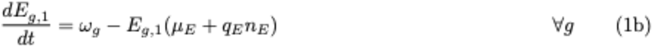

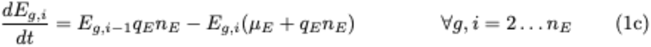

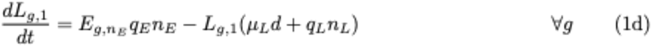

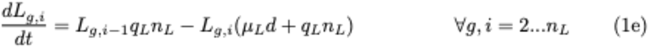

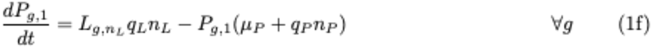

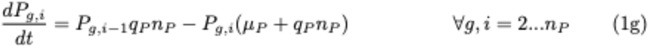

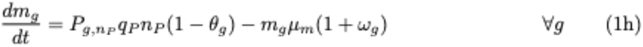

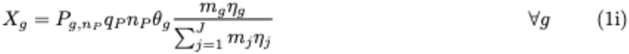

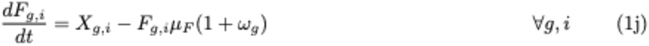

Each geographic node contains equations of motion (Eq.1a-1j) calculating oviposition (κ), the presence of juvenile life stages egg (*E*), larva (*L*), and pupa (*P*), and adult life stages male (*m*) and female (*F*). Adults are assumed to mate a single time; variable *X* represents mated females. The organisms within a shared life stage and node are parameterized by the same death (*μ*) and maturation (*q*) rates.^50^ Larval stage mortality is additionally regulated by logistic density dependence (*d* = 1 + ∑(*L*)/*k*, where *k* represents carrying capacity). Following multiple examples in ecology and public health applications, we use Erlang distributions (*n*)^51–53^ to incorporate flexible dwell times helpful for modeling the environmentally-influenced life stage duration of juvenile mosquitoes.

Life stage variables are indexed by genotype (*g*) to reflect a particular pattern of inheritance and Erlang substage (*i*). All variables are implicitly indexed by time; for brevity of presentation we omit these in Eq. 1. In the case study examples presented by this work, *g* differentiates between RIDL-modified and wildtype individuals, while mortality rates *μ* and development rates *q* are dynamically calculated for each life stage using the *Ae. aegypti*-specific temperature-sensitive functional forms developed by Rossi et. al. (2014).^54^ Fitness cost of the RIDL technology is parameterized with mid-range estimates from field trial data,^14^ with transgenic males penalized by a 3.1% decrease in mating fitness (*η_g_*) and an 18% additional increase (*ω_g_*) in mortality.

Parameter *β_g_* represents female fecundity, while *α_g_* represents male fecundity. *Г_g_* and *T^g^* are genotype- specific inheritance and survival probability, respectively, and *θ_g_* dictates the male to female offspring emergence ratio. Values for *Г_g_* used in the worked examples are from Sánchez et. al. (2020)^55^; the formulation of Eq.(1a) and (1i) follow Sánchez et. al. (2020).^56^ The ODE system is initialized at equilibrium by setting the left hand side to zero and solving (see Methods Section 5: The Julia advantage).

While the data model establishes an experimental record of the parameters used to initialize a dynamic simulation in GeneDrive.jl, the solver permits updates to these values over the course of a dynamic model run (see Methods Section 5: The Julia advantage). This feature facilitates investigating the effect of system perturbations. It is enabled by the information flow that characterizes the dynamic model (Methods Figure 2). In the examples shown, a use case is the introduction of RIDL-modified organisms into the modelled population at the various timesteps dictated by different optimized policies (see Results). Other work has applied this capability to alter the environmental conditions to which simulated organisms are responding.^29^

**Methods Figure 2:**
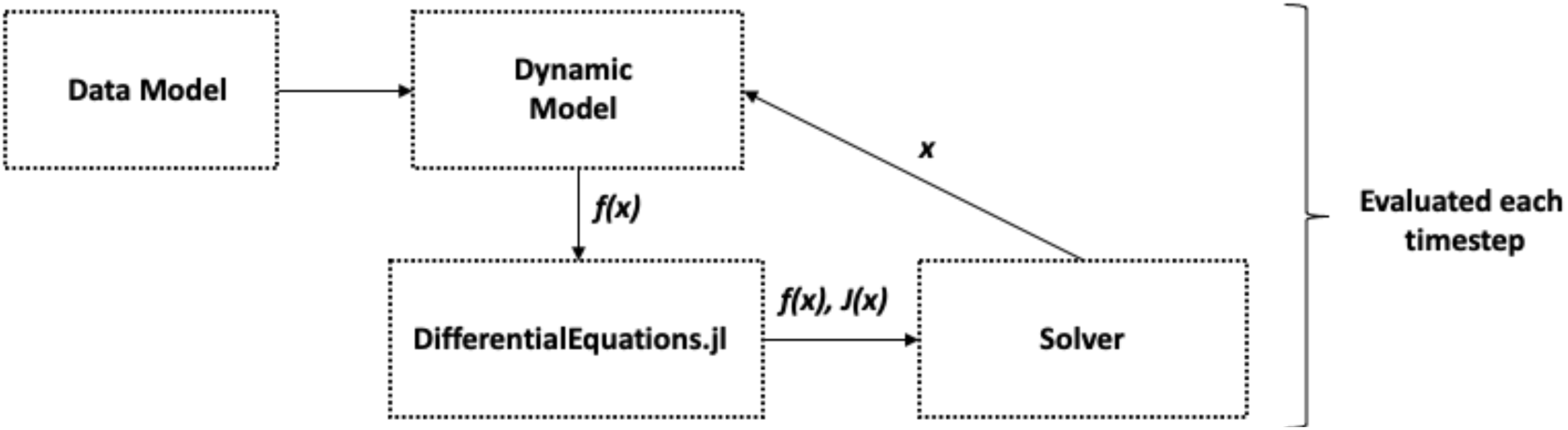
Dynamic model information flow.

The information flow key to such computational experiments is enabled by modular design (see Methods Section 5: The Julia advantage). Modularity also enforces the separation of simulation and solving algorithms, curtailing the need for extensive code rewrites when models are updated and run. For example, to create the dynamic model once the data model is built, all that is required is solver selection followed by simulation using the ‘solve_dynamic_model’ API (Code Block 2). Outputs may then be pre- processed for analysis using ‘format_dynamic_model_results.’ Modular design likewise permits solution methods to be easily exchanged, allowing the exploration of alternative backend solvers to improve performance. The dynamic model examples shown as case studies use Tsit5, an explicit Runge-Kutta method recommended for most non-stiff systems.^32^ However, there are a robust suite of options compatible with or included in the DifferentialEquations.jl platform upon which the GeneDrive.jl dynamic model is built (see Methods Section 5: The Julia advantage).

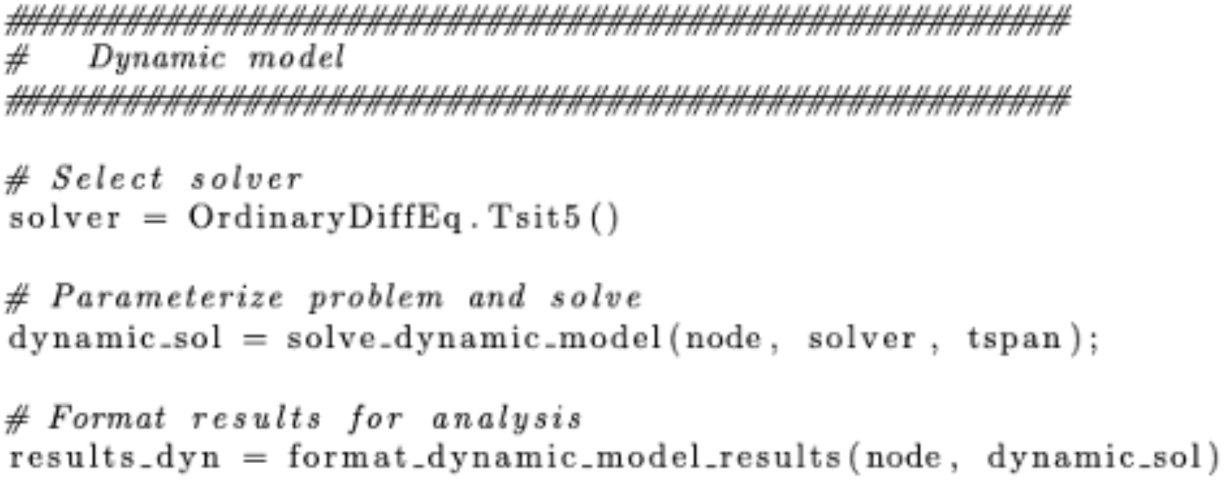
***Code Block 2:*** *Creating and solving the dynamic model, then formatting outputs for further analysis, is straightforward in GeneDrive.jl*.

### Section 4: Decision model

The GeneDrive.jl decision model is a nonlinear program (NLP) formulated by discretizing the dynamic model’s system of ODEs using Euler approximations with a daily timestep. It determines the optimal set of control (also called decision) variable values in pursuit of a given objective (goal). The objective is a function that maximizes or minimizes, and can be a measure of performance, cost, or any other metric of interest. The “best” set of decision variables to satisfy an objective function are often referred to as the optimal schedule or optimal policy and are subject to constraints. Constraints, or restrictions on the feasible solution space, are defined using mathematical equalities or inequalities.

The equality constraints of the GeneDrive.jl optimization problem are the discretized system of ODEs representing the vector population. Thus, they inform the feasibility of a given simulation with biological details. Constraints representing non-biological or operational limitations, such as resource availability, enter the decision model as inequality constraints. These are assigned on a per-node basis using the data model, enabling the exploration of spatially differentiated policy combinations. Like the dynamic model, decision model parameters are species-specific and populated by data model values. Unlike the dynamic model, solution methods evaluate problem information over the full time horizon rather than at each timestep (Methods Figure 3).

**Methods Figure 3:**
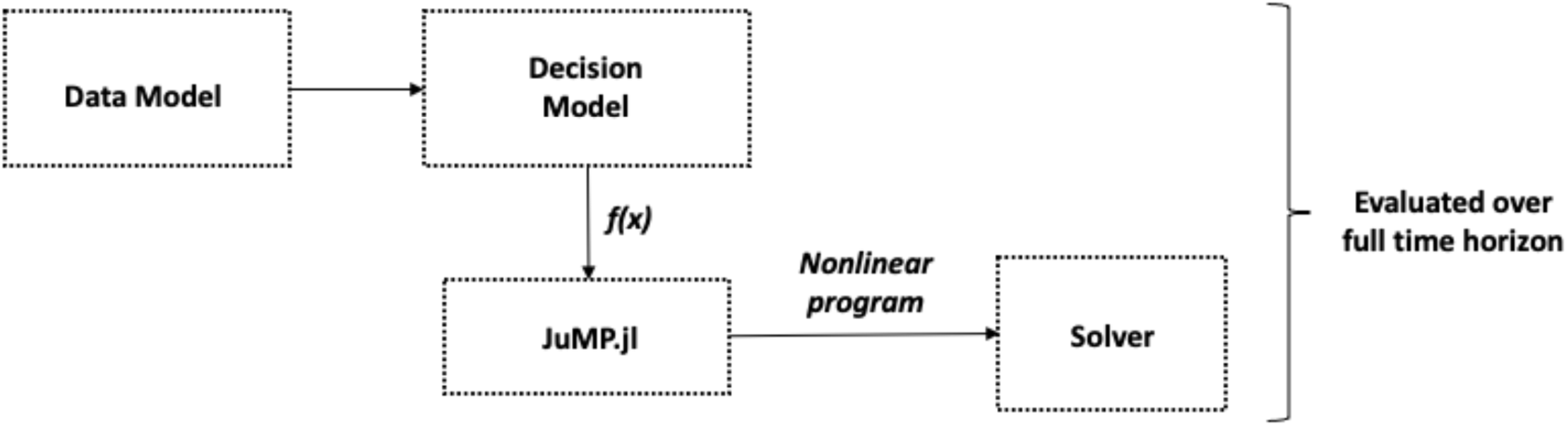
Decision Model information flow.

The default nonlinear solver in GeneDrive.jl is the Interior Point OPTimizer (Ipopt),^57^ a free software library for largescale optimization. This choice may be updated by users interested in exploring different solvers or augmenting the Ipopt optimizer with additional algorithms to accelerate solutions or improve accuracy. Ipopt internally utilizes a linear solver. Various free and commercial linear solvers can be chosen from among a list in the Ipopt online documentation^57^ that are compatible with the JuMP.jl platform used to create the GeneDrive.jl decision model (see Methods Section 5: The Julia advantage). While sample objective functions are supplied in the GeneDrive.jl package, optimization goals are unique to package users and thus also customizable.

Again, modularity simplifies both problem specification and simulation, maximizing code re-use. After populating the ‘ReleaseStrategy’ structure with context-specific operational constraints and assigning those values to the ‘Nodè structure of interest in the data model, a solving algorithm is selected. The deterministic optimization problem is then parameterized using ‘create_decision_model’, solved using solve_decision_model_scenarios, and results formatted for analysis with ‘format_dynamic_model_results’ (Code Block 3, left panel). The stochastic optimization model is differentiated via the form of temperature structure inputs to the updated data model: it takes a matrix of time series together with associated probabilities, unlike the single time series used as the input to the deterministic optimization (Code Block 3, right panel). Operational constraints specific to the experiment of interest in this case study permit only male organisms modified using the RIDL technology to be released, with the timing of those deployments allowed weekly in batches numbering 50,000 organisms or less at each release.

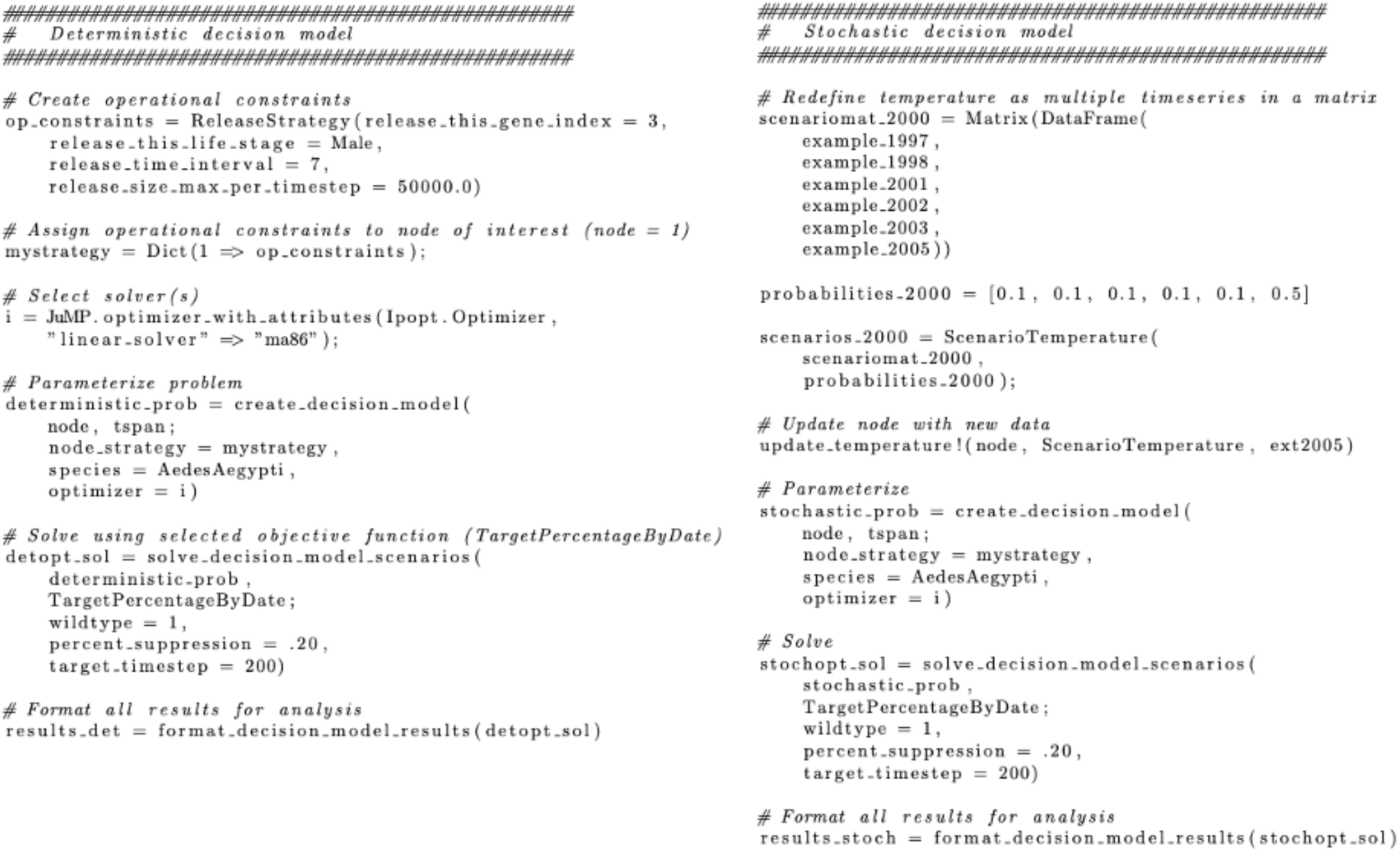
***Code Block 3:*** *Creating the deterministic and stochastic optimization models*.

### Section 5: The Julia advantage

The entire GeneDrive.jl stack is free and open source, lowering barriers to usage and enhancing the transparency of its methods and outputs. This is enabled by the Julia scientific programming language, as well as the packages within that ecosystem used to develop this work. The equilibrium of the dynamic and decision model systems is found using NLSolve.jl,^58^ a library for solving nonlinear equations. The dynamic model itself is built upon DifferentialEquations.jl,^32^ a suite for numerically solving differential equations that also furnishes efficient differential equation solution methods. It is DifferentialEquations.jl that allows the use of callback functions within GeneDrive.jl. This means that custom changes can be made to data values or functions at specific points during the execution of a simulation, permitting users to model exogenous inputs to the system including the addition of new species and the manifestation of heatwaves. The decision model is developed using JuMP.jl,^33,59^ a domain specific modelling language (DSL) and collection of supporting packages for mathematical optimization that is embedded in Julia.

Finally, Julia supports a growing community of developers interested in biological questions and expressly focused on cutting edge advances in scientific programming.^31,60^

## Code and data accessibility

The GeneDrive.jl software is located on GitHub at https://github.com/vnvasquez/GeneDrive.jl; all code and data inputs required to reproduce the work in this paper is included in the https://github.com/vnvasquez/paper-genedrivesoftware repository. Model dynamics are validated via tests in that repository as well as against empirical study results in Vásquez et. al. (2023).

